# Immunoinformatics Prediction of Epitope Based Peptide Vaccine Against *Madurella mycetomatis Translationally Controlled Tumor Protein*

**DOI:** 10.1101/441881

**Authors:** Samira Munir Bolis, Walaa Abdullah Omer, Mohamed Anwar Abdelhamed, Masajed Abdelmagid Shambal, Esameldeen Ahmed Adam, Mohammed Abaker Abass, Wiaam Abdelwahab Abdalla, Suzan Hashim Is-haq, Aisha Abubakralsiddig Abdalla, Abeer Algaali Zeinalabedeen, Omar Hashim Ahmed, Mohamed A Hassan

## Abstract

**Background:** *Madurella. mycetomatis* is most common causative agent of mycetoma in Sudan and worldwide. No vaccines are available till now so design of effective vaccine is essential as protection tool. Peptide vaccine can overcome the common side effects of the conventional vaccines. The aim of this study was to design peptide based vaccine for *M.Mycetomatis Translationally Controlled Tumor Protein (TCTP)* using immunoinformatics tools.

**Materials and methods:** *TCTP* sequences were retrieved from NCBI and then processed using BioEdit program to determine conserved regions and different immunoinformatics tools from IEDB. Population coverage analysis was performed for the most promising epitopes. Homology modelling was performed to show their structural positions in *TCTP.* Protein analysis was done using Expasy (ProtParamsotware).

**Results and conclusion:** Four epitopes passed the Bepipred, Emini, Kolaskar and Tongaonkar tools. 111 epitopes were predicted to interact with MHCI alleles with IC50 < 500 nM, three of them were most promising. 274 predicted epitopes were interacted with MHCII alleles with IC50 < 100 nM, four of them were most promising. The epitope (YMKSVKKAL) was the most promising one concerning its binding with MHCI alleles, while (FRLQSTSFD) was the most promising for MHC II. The epitope (YLKAYMKSV) is shared betweenMHC I and II. For the population coverage of *M. Mycetomatis TCTP* vaccine Sudan (90.39%) had the highest percentage for MHC I. This is the first computational vaccinology study conducted in mycetoma caused by *M. Mycetomatis* using *TCTP.*

## Introduction

Mycetoma is a granulomatous subcutaneous chronic progressive infection. It is characterized by a painless swelling, development of draining sinuses that discharge different colors of pus and grains (1–3). Mycetoma is subdivided into actinomycetoma and eumycetoma (1, 4, 5).. It is distributed worldwide, but it is endemic in tropical and subtropical regions (6–8). These region sknown as the “Mycetoma belt” including (Sudan, Somalia, Yemen, Senegal, India, Mexico, Venezuela, Colombia and Argentina) while the most reported cases were from Sudan and Mexico, Sudan being the most endemic country (9) (10, 11). Mycetoma can be found in areas geographically close proximity to Tropic of Cancer (1). It is categorized as one of the neglected tropical diseases at 2013 (6, 12). Accurate data on its incidence and prevalence are not available (12). The most common causative organism of mycetoma in Sudan and worldwide is *Madurella mycetomatis* (13–15). Mycetoma is still widely distributed in Sudan with no effective treatment, no vaccines are available till now hence the design of an effective peptide vaccine is essential as protection tool against the disease (12). This study aimed to predict an insilico peptide based vaccine for mycetoma caused by *M. mycetomatis* using immunoinformatics.

The defense mechanisms against fungi usually range from early non-specific immune response to activation of specific adaptive immune responses by the production of cytokines of T helper-1 and T helper-2. Cell-mediated immunity can play a role in the pathogenesis of eumycetoma (16). The first well-characterized immunogenic antigen is *Translationally Controlled Tumor Protein (TCTP)*. Significant IgG and IgM immune responses against this protein were determined. The level of antibodies is directly proportional to lesion size and duration of the disease (17). Different laboratory-based diagnostic techniques and tools were developed to determine and identify the causative agents of mycetoma (18). Early detection and treatment are important to reduce morbidity and improve treatment outcomes (5, 16). Antifungals are used for the treatment of eumycetoma, amputations and recurrences are common (4, 8). Recently understanding of complex interactions between fungi and host led to exploration and design of novel vaccines (19). The first genome sequence for this strain in Sudan was presented in 2016, better therapies for mycetoma will be developed using this sequence (13). *TCTP* is abundantly expressed in wide range of eukaryotic organisms located in the cytoplasm and the nucleus (20, 21). It is highly conserved protein and its expression is associated with malignancy and chemoresistance. *TCTP* plays important role in physiological events, such as immune responses ,cell proliferation and cell death (20). *TCTP* is simultaneously the first identified monomolecular vaccine candidate for *M.mycetomatis* (17). Next to the *TCTP*, two antigenic proteins of *M.mycetomatis* were discovored, the glycolytic enzymes *Fructose Biphosphate Aldolase* (*FBA*) and *Pyruvate Kinase (PK)*, these antigens might also be useful as vaccine-candidates in the prevention of mycetoma (22). The gene for this protein was found to be present in two variants in *M. mycetomatis,* with difference in 13% amino acids between the two encoded proteins (17). This is the first computational vaccinology study conducted in mycetoma caused by *M.mycetomatis* using *TCTP*.

## Materials and methods

### Protein sequence retrieval

Nine *Madurella mycetomatis TCTP* sequences were retrieved from variants one and two from the National Center for Biotechnology Information (NCBI) (https://www.ncbi.nlm.nih.gov/protein/?term=mycetoma+and+TCTP) database in August 2017.

### Determination of conserved regions

Multiple sequence alignment (MSA) using ClustalW as in the BioEdit program, version 7.2.5 was used to obtain conserved regions among *Madurella. mycetomatis TCTP* (23).

### B-cell epitope prediction

The reference *Madurella. mycetomatis TCTP* sequence was subjected to Bepipred linear epitope, Emini surface accessibility and Kolaskar and Tongaonkar antigenicity methods in Immune Epitope Data Base (IEDB) (22–24) (http://www.iedb.org/), that predict the probability of specific regions in the protein to bind to B cell receptor, being linear, in the surface and immunogenic, respectively.

### Prediction of linear B-cell epitopes

Bepipred from immune epitope database (http://tools.iedb.org/bcell/) was used as linear B-cell epitopes prediction tool from the conserved region with a default threshold value of 0.240(25).

### Prediction of surface accessibility

By using Emini surface accessibility prediction tool of the (IEDB) (22–24) (http://tools.iedb.org/bcell/result/). The surface accessible epitopes were predicted from the conserved region holding the default threshold value 1.000 (26).

### Prediction of epitopes antigenicity sites

The kolaskar and tongaonker antigenicity method (http://tools.iedb.org/bcell/result) was used to determine the antigenic sites with a default threshold value of 1.024 (27).

### MHC class I binding predictions

Analysis of peptide binding to MHC class I molecules was assessed by the (IEDB) MHC I prediction tool at (http://tools.iedb.org/mhci/result/). Peptide complex presentations to T lymphocytes undergo several steps. The attachment of cleaved peptides to MHC molecules step was predicted. Prediction methods was achieved by Artificial Neural Network (ANN) method (28–32). Prior to prediction, all epitope lengths were set as 9amino acids, all conserved epitopes that bind to alleles at score less than 500 inhibitory concentration (IC50) were selected for further analysis (32).

### MHC class II binding predictions

Analysis of peptide binding to MHC class II molecules was assessed by the IEDB MHC II prediction tool at (http://tools.iedb.org/mhcii/result/) (33, 34). For MHC-II binding predication, human allele references set were used., NN-align uses the artificial neural networks that allows for simultaneous identification of the MHC class II binding core epitopes and binding affinity from IEDB was used as prediction methods for MHC II (35). All conserved epitopes that bind to many alleles at score less than 100 half-maximal inhibitory concentration(IC50) were selected for further analysis (33).

### USCF Chimera version 1.8

It is a program for visualization and analysis of molecular structures and related data with high-quality images and animations. It is produced by University of California, San Francisco. Fallowing step by step to generate models for *Madurella mycetomatis TCTP*. (Available at https://www.cgl.ucsf.edu/chimera/).

### Population coverage calculation

Selected MHC-I and MHC-II interacted alleles by the IEDB population coverage calculation tool at (http://tools.iedb.org/tools/population/iedb_input) (36).

## Results

### Protein sequence retrieval

Retrieved *TCTP* sequences and their accession numbers are: ABB20812, ABB20815, ABB20814, ABB20813,ABB20811, ABB20810, ABB20809, ABB20808, ABB20807.

### Alignment

*Madurella mycetomatis TCTP* conserved regions are shown in (figure 1).

### Prediction of B-cell epitope

In Bepipred Linear Epitope Prediction method; the average binders score of *TCTP* to B cell was 0.240, with maximum score 1.974 and a minimum score −1.296 (figure 2). Seven epitopes were predicted eliciting B lymphocyte from the conserved regions and all values equal or greater than the default threshold 0.24.

In Emini surface accessibility prediction; the average surface accessibility areas of the protein was scored as 1.000, with a maximum of 2.963 and a minimum of 0.073(figure 3). 37 epitopes were potentially in the surface by passing the default threshold 1.000.

In Kolaskar and Tongaonkar antigenicity; the average of the antigenicity was 1.024, with a maximum of 1.188 and minimum of 0.892(figure 4). 15 epitopes gave score above the default threshold 1.024.

There are four epitopes successfully passed the three tools (DEVKEFETKAQAYV, EVKEFETKAQAYV, EFETKAQAYV, ETKAQAYV) as shown in (table 1).

The positions of proposed conserved B cell epitopes in structural level of *TCTP* of *Madurella. mycetomatis* are shown in (figure 5).

### Prediction of cytotoxic T-lymphocyte epitopes and interaction with MHC class I

The reference *TCTP* sequence was analyzed using (IEDB) MHC-1 binding prediction tool to predict T cell epitope suggested interacting with different types of MHC Class I alleles, based on Artificial Neural Network (ANN) with half-maximal inhibitory concentration (IC50) <500 nm.111peptides were predicted to interact with different MHC-1alleles. The most promising epitopes and their corresponding MHC-1 alleles are shown in (Table 2). Their positions at the structural level were determined by the aid of the chimera software as shown in (Figures6,Figures7).

### Prediction of The T cell epitopes and interaction with MHC class II

Reference *TCTP* variant was analyzed using (IEDB) MHC-II binding prediction tool based on NN-align with half-maximal inhibitory concentration (IC50) <100 nm; there were 274 predicted epitopes found to interact with MHC-II alleles.

The results of top four epitopes are listed in (Table 3). Their positions at the structural level were determined by the aid of the chimera software as shown in (Figure 8,Figure 9).

### Protein analysis

*M.mycetomatis TCTP* protein sequence was analyzed using both Bioedit program, version 7.0.9.0 and Expasy (ProtParam software) (https://web.expasy.org/cgi-bin/protparam/protparam) softwares′ tools (figure 10). Number of amino acids: 141. Molecular weight: 15794.80

Total number of negatively charged residues (Asp + Glu): 25. Total number of positively charged residues (Arg + Lys): 17. The Atomic composition of *Madurella mycetomatisTCTP* is shown in (Table 4).

Formula: C705H1095N177O224S5. Total number of atoms: 2206. The N-terminal of the sequence considered is V (Val). The estimated half-life is: 100 hours (mammalian reticulocytes, in vitro), >20 hours (yeast, in vivo) and >10 hours (Escherichia coli, in vivo).

Instability index: The instability index (II) is computed to be 29.38 (the protein is stable). Grand average of hydropathicity (GRAVY): −0.428

### Population coverage calculation

All promising MHC I and MHC II binders of *Madurella mycetomatis TCTP* (epitopes with high binding affinity with different sets of alleles) were assessed for population coverage against the whole world and Sudan (table 5).

**Fig 1.**
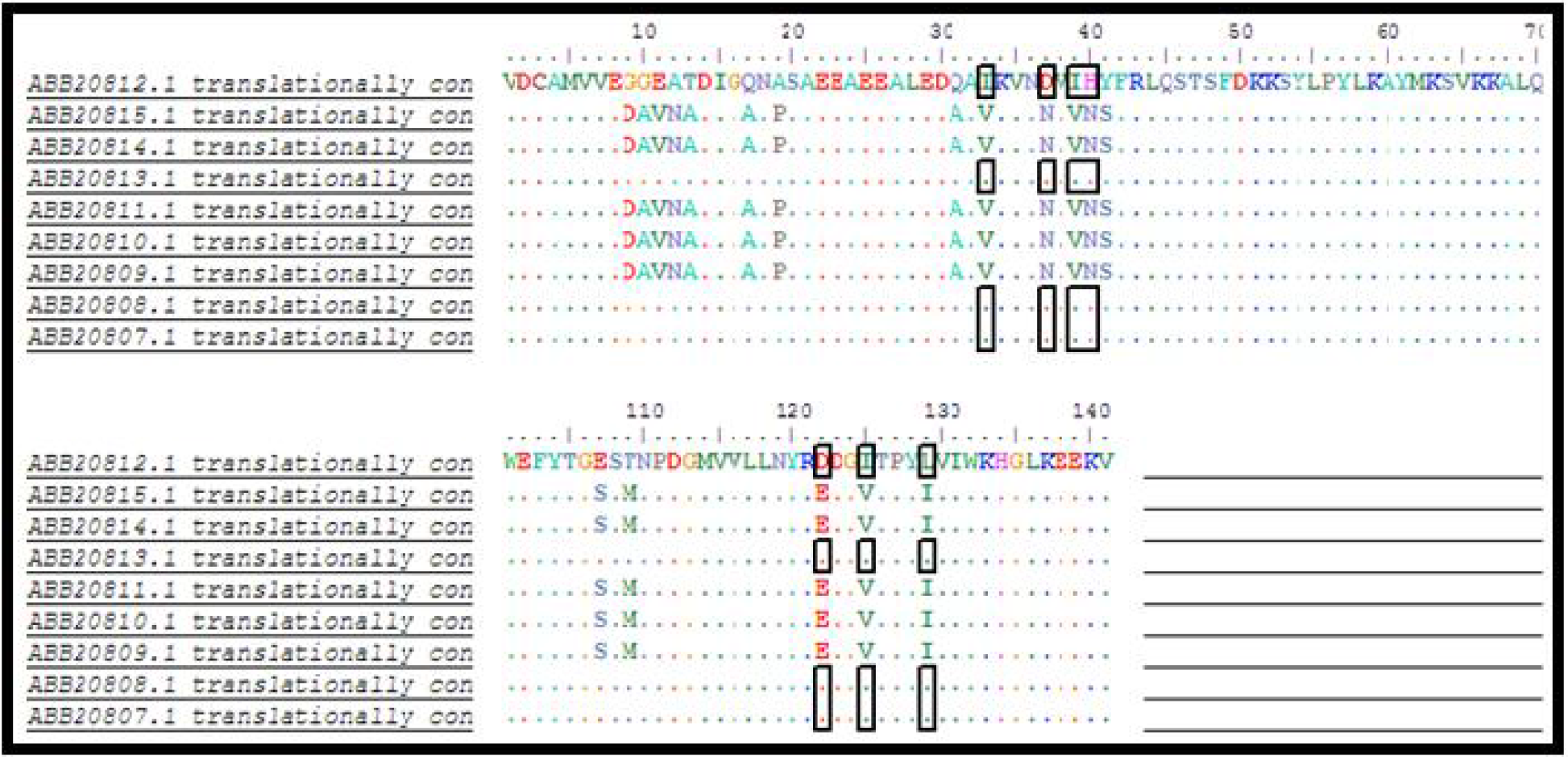
Multiple sequence alignment, dots show the conservancy between sequences *The alignment is done using BioEdit tool.

**Fig 2.**
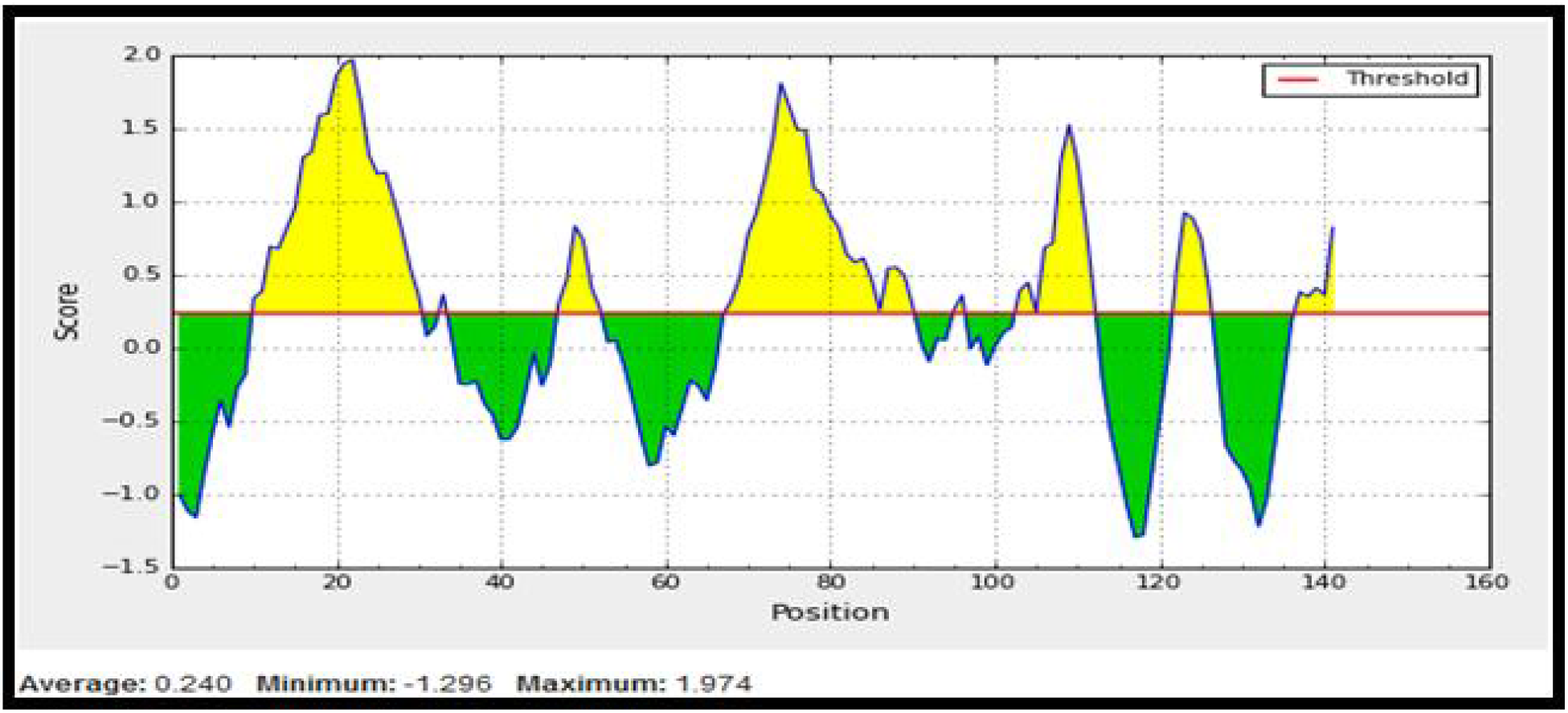
Bepipred linear epitope prediction. Yellow areas above threshold (red line) are proposed to be a part of B cell epitope. While green areas are not

**Fig 3.**
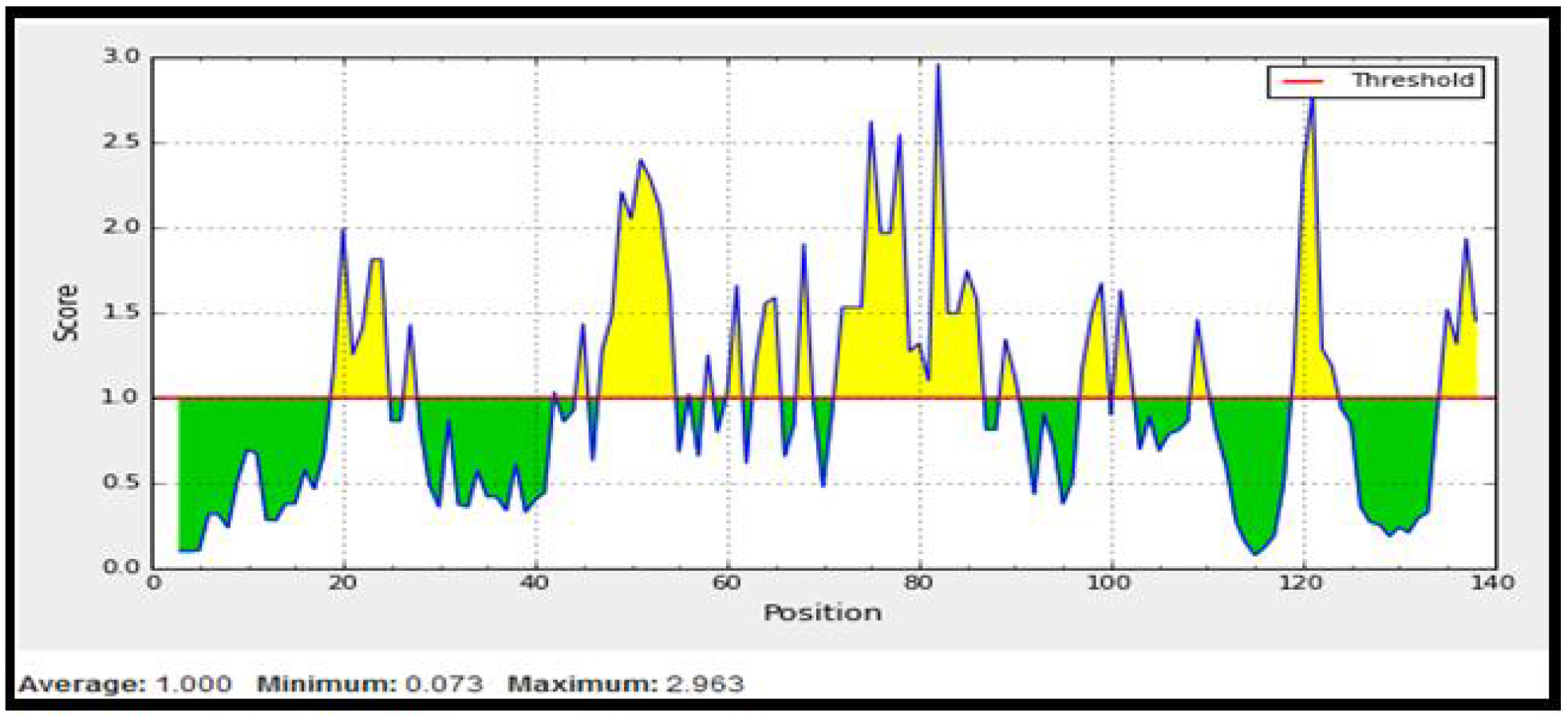
Emini surface accessibility prediction. Yellow areas above threshold (red line) are proposed to be a part of B cell epitope. While green areas are not.

**Fig 4.**
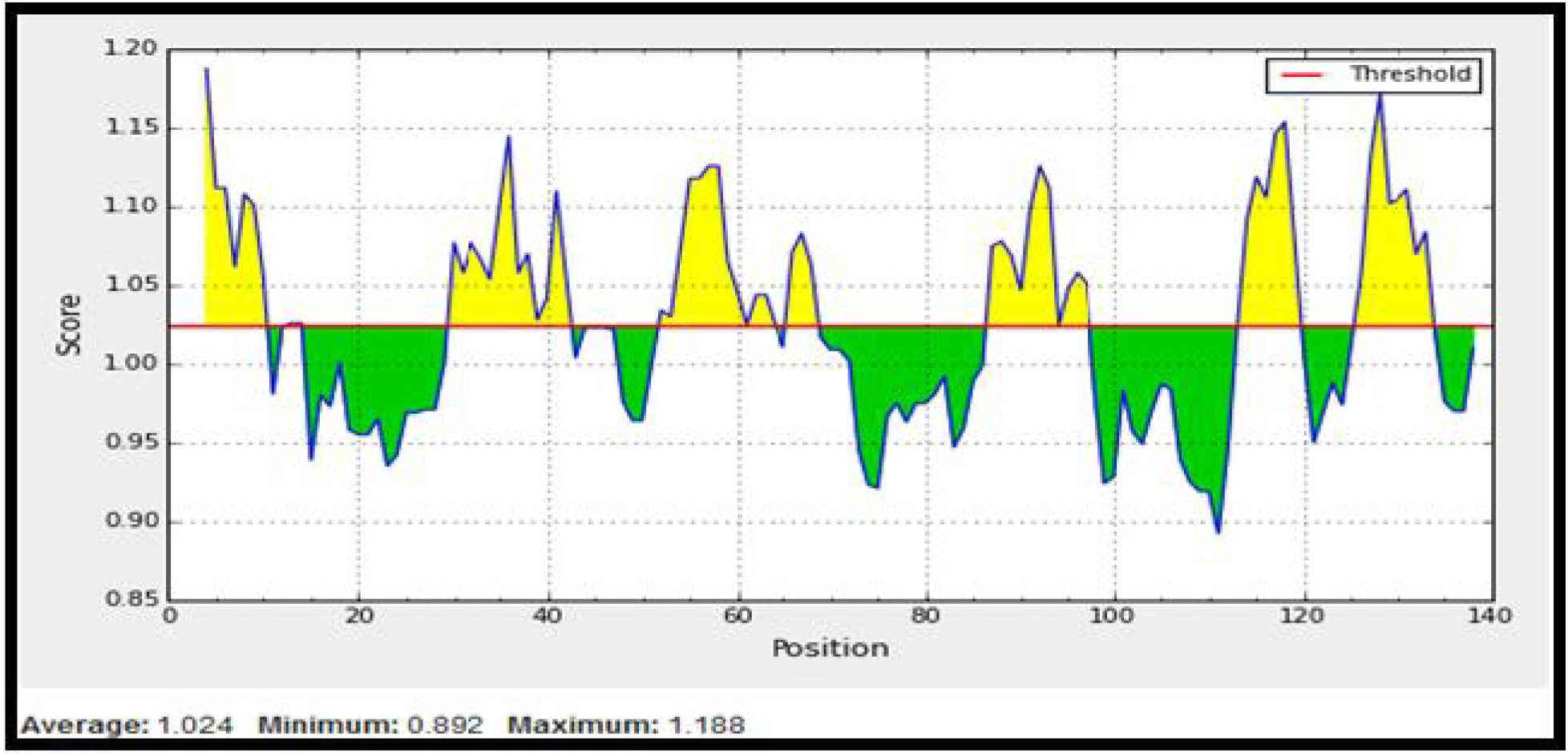
Kolaskar and Tongaonkar antigenicity prediction. Yellow areas above threshold (red line) are proposed to be a part of B cell epitope. While green areas are not.

**Table 1:**
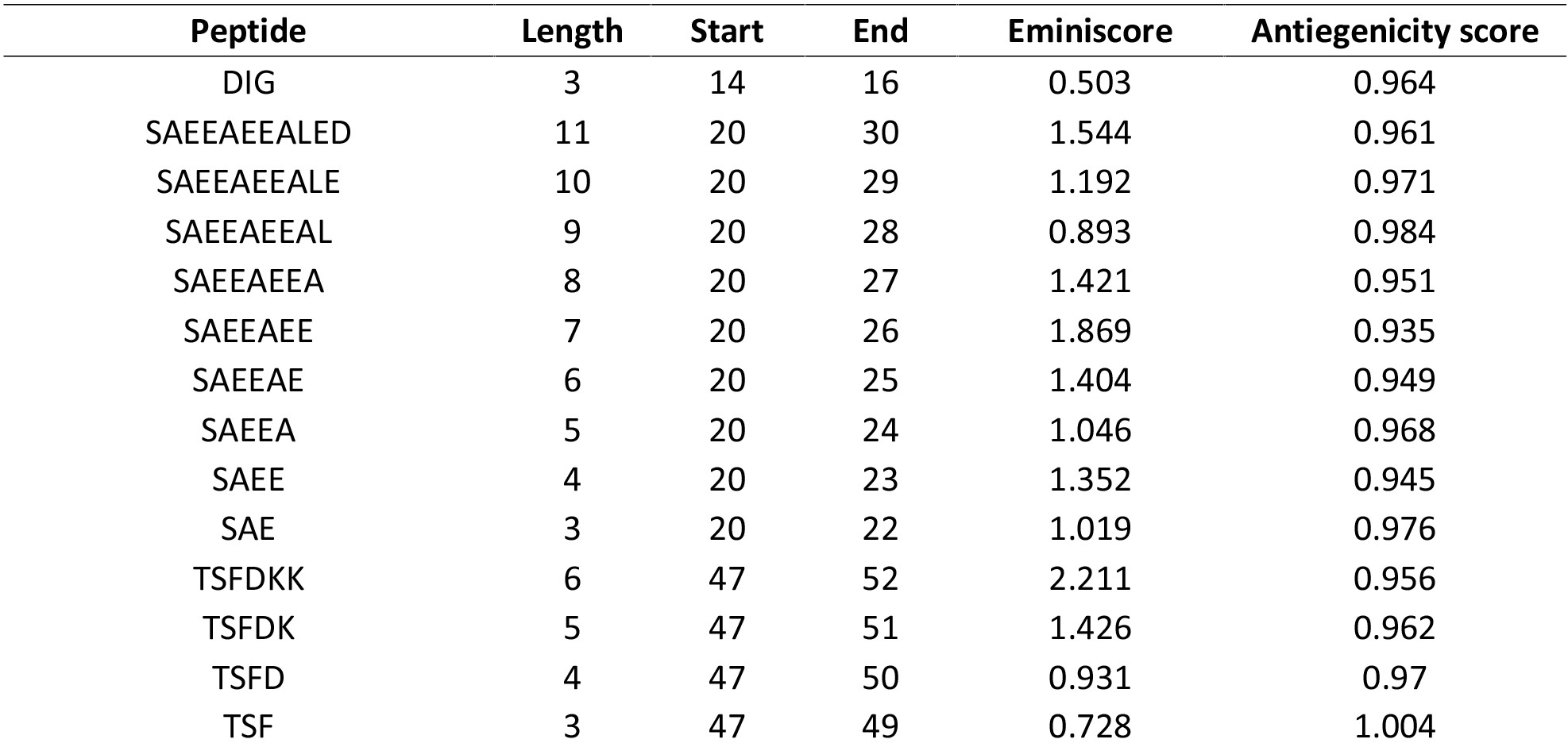
B cell epitopes prediction, Peptides in the red color have passed the three tests of the B-cell

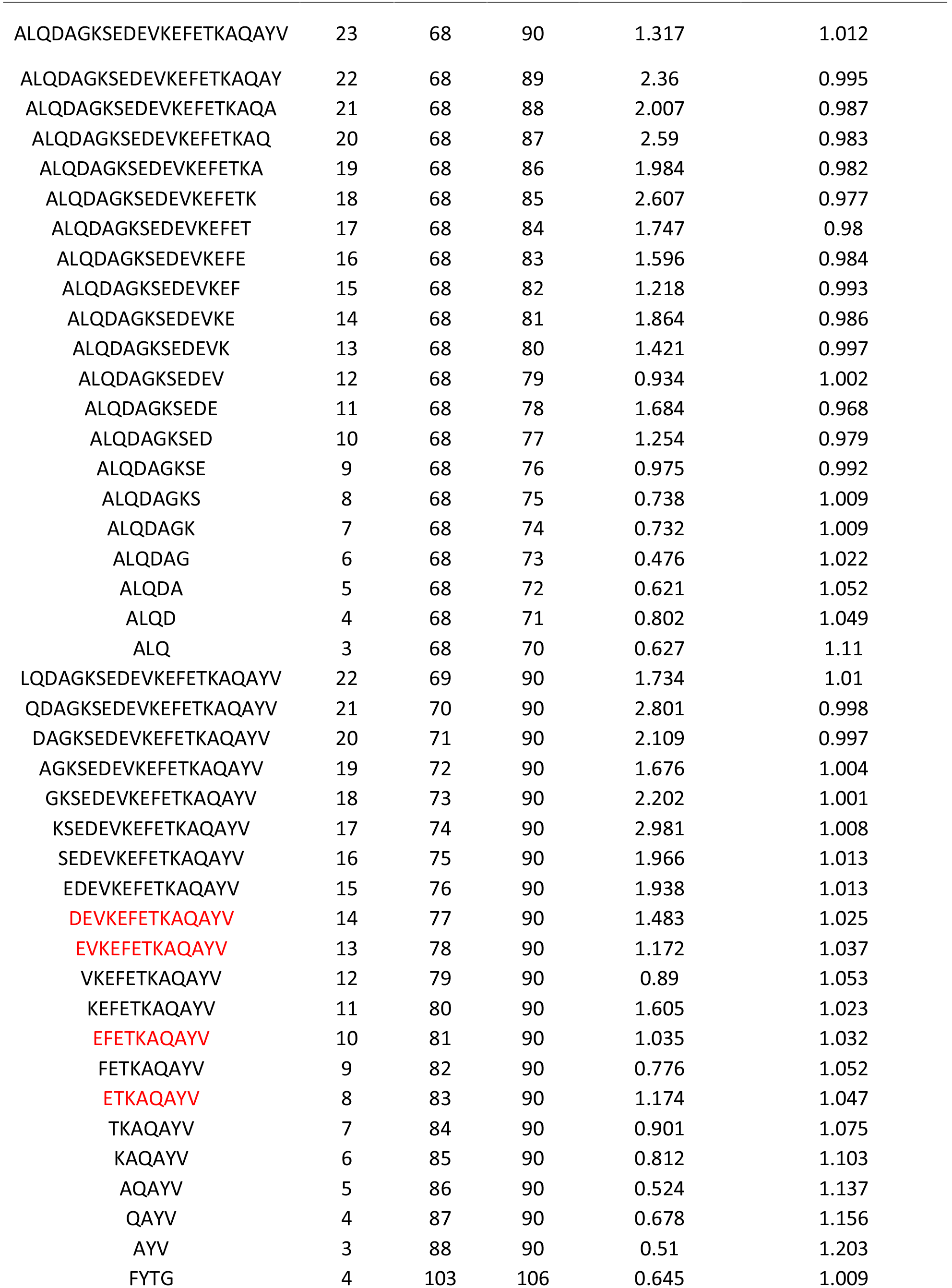

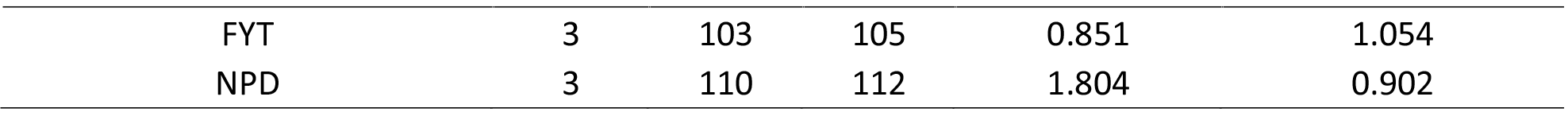

**Fig 5.**
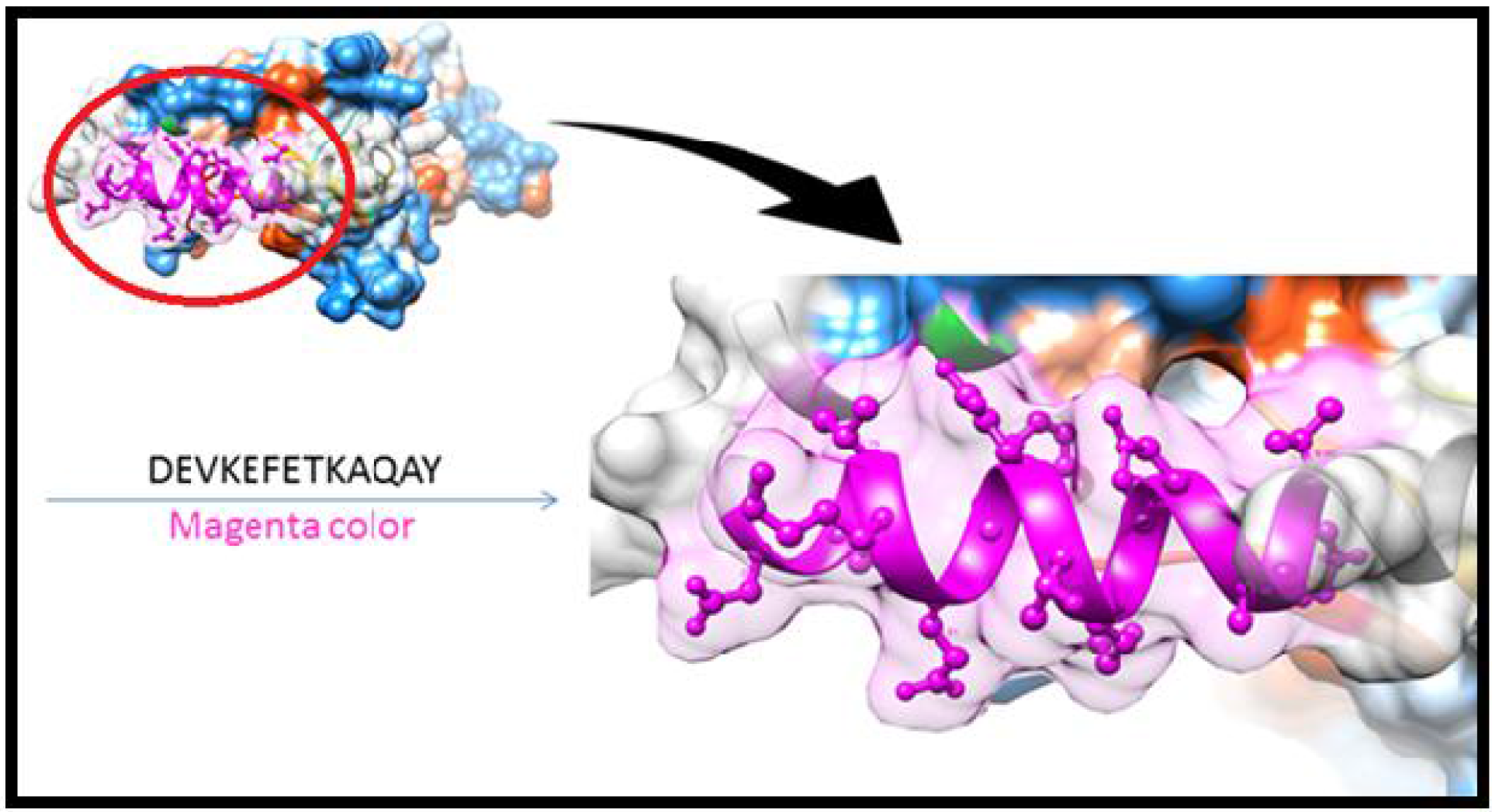
B cell epitopes proposed. the arrow show positions of (DEVKEFETKAQAY) with magenta color in structural level of *Madurella. Mycetomatis TCTP.*

**Table 2.**
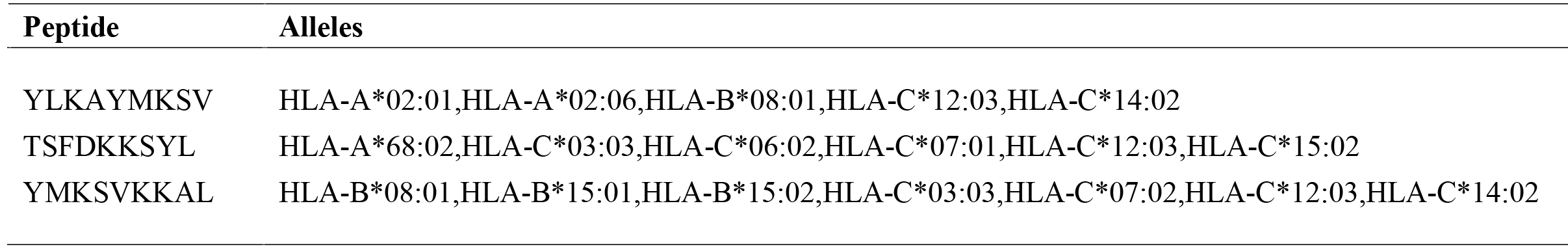
The most Promising epitopesand their corresponding MHC-1 alleles

**Fig 6.**
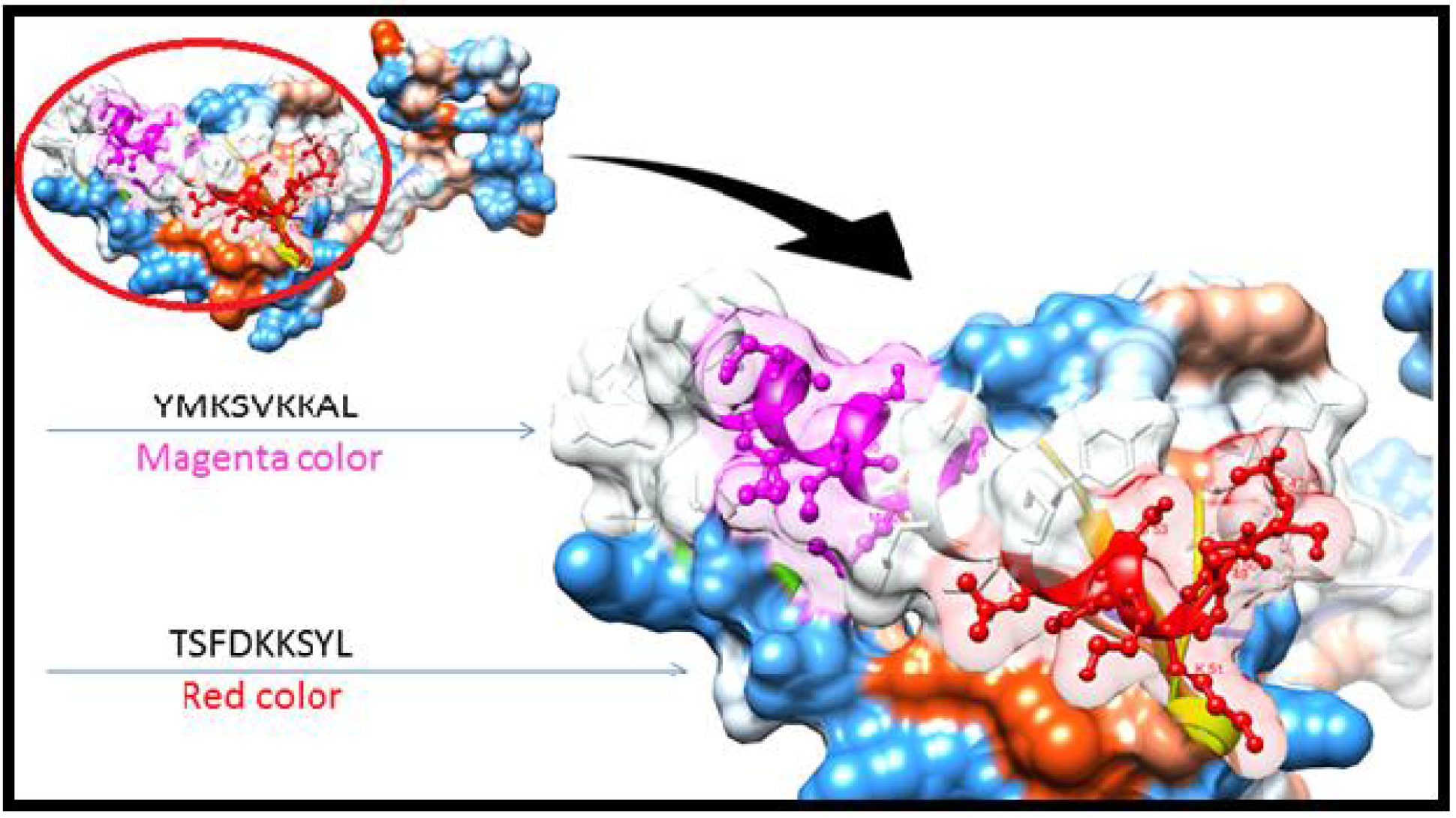
T cell epitopes proposed that interact with MHCI. The arrows show position of (YMKSVKKAL)with magenta color, (TSFDKKSYL) with red color in structural level of *Madurella. Mycetomatis TCTP*

**Fig 7.**
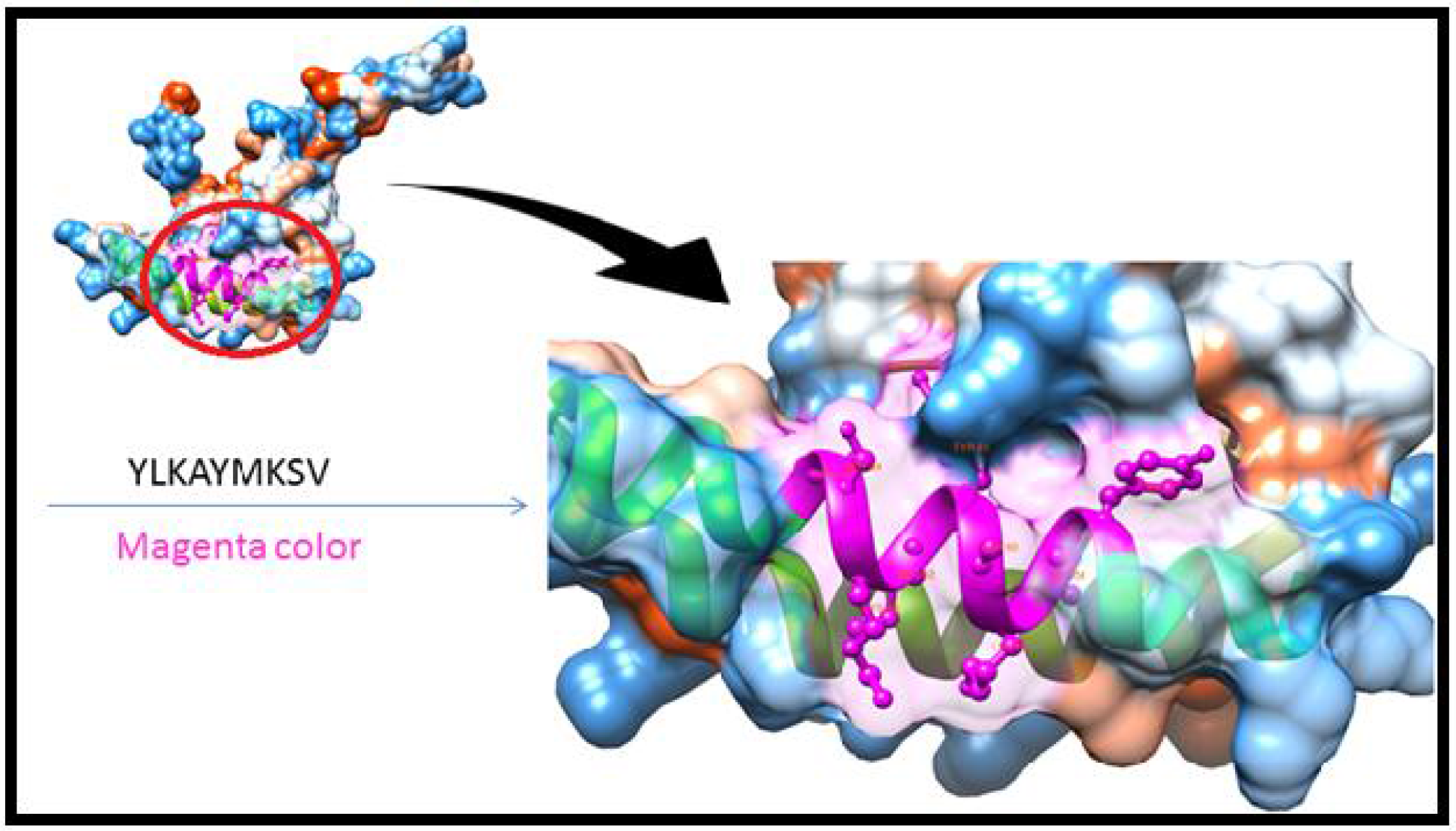
cell epitopes proposed that interact with MHCI. The arrow show position of (YLKAYMKSV)with magenta color in structural level of *Madurella Mycetomatis TCTP*

**Table 3.**
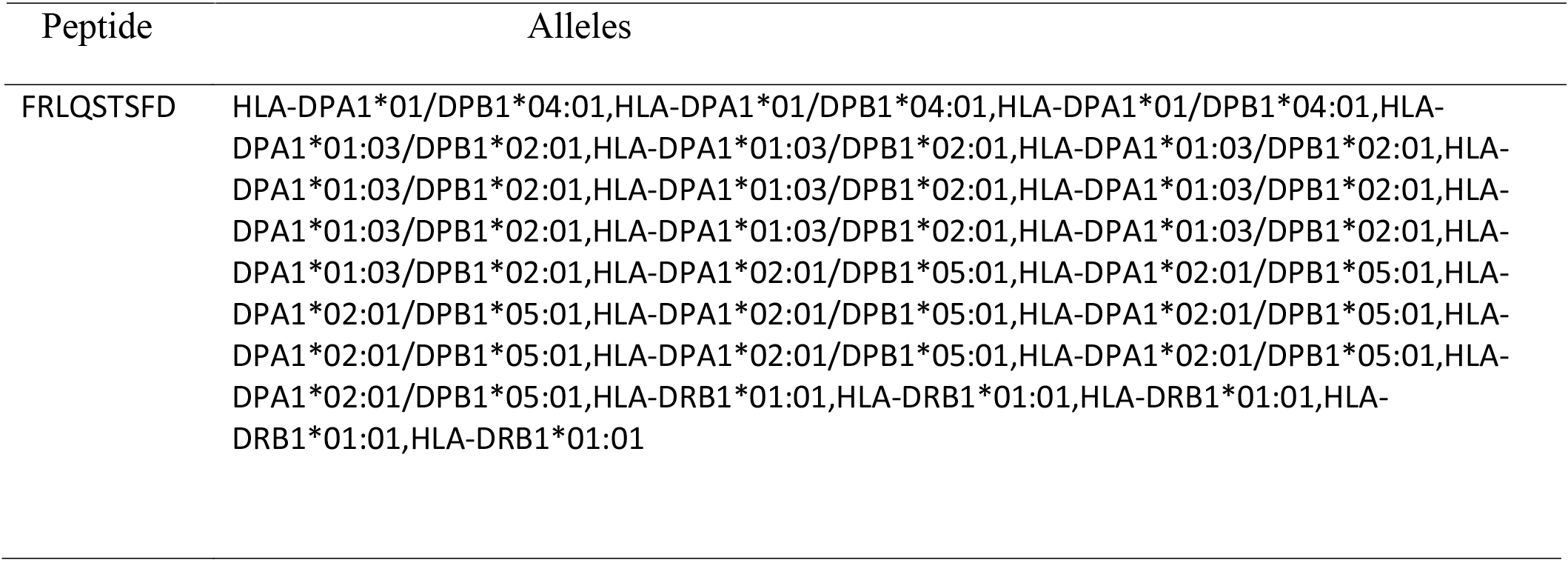
The most Promising epitopes and their corresponding MHC-**II** alleles

|  |  |
| --- | --- |
| YLKAYMKS | HLA-DRB1*11:01,HLA-DRB1*11:01,HLA-DRB1*11:01,HLA-DRB1*11:01,HLA-DRB1*11:01,HLA-DRB1*11:01,HLA-DRB1*11:01,HLA-DRB1*11:01,HLA-DRB1*11:01,HLA-DRB1*11:01,HLA-DRB1*15:01,HLA-DRB1*15:01,HLA-DRB1*15:01,HLA-DRB1*15:01,HLA-DRB1*15:01,HLA-DRB1*15:01,HLA-DRB1*15:01,HLA-DRB1*15:01 |
| YLPYLKAYM | HLA-DRB1*15:01,HLA-DRB3*01:01,HLA-DRB3*01:01,HLA-DRB3*01:01,HLA-DRB3*01:01,HLA-DRB5*01:01,HLA-DRB5*01:01,HLA-DRB5*01:01,HLA-DRB5*01:01,HLA-DRB5*01:01,HLA-DRB5*01:01,HLA-DRB5*01:01 |
| LQSTSFDKK | HLA-DRB1*04:05,HLA-DRB1*04:05,HLA-DRB1*07:01,HLA-DRB1*07:01,HLA-DRB1*07:01,HLA-DRB1*07:01,HLA-DRB1*07:01,HLA-DRB1*07:01,HLA-DRB1*07:01,HLA-DRB1*08:02 |

**Fig 8.**
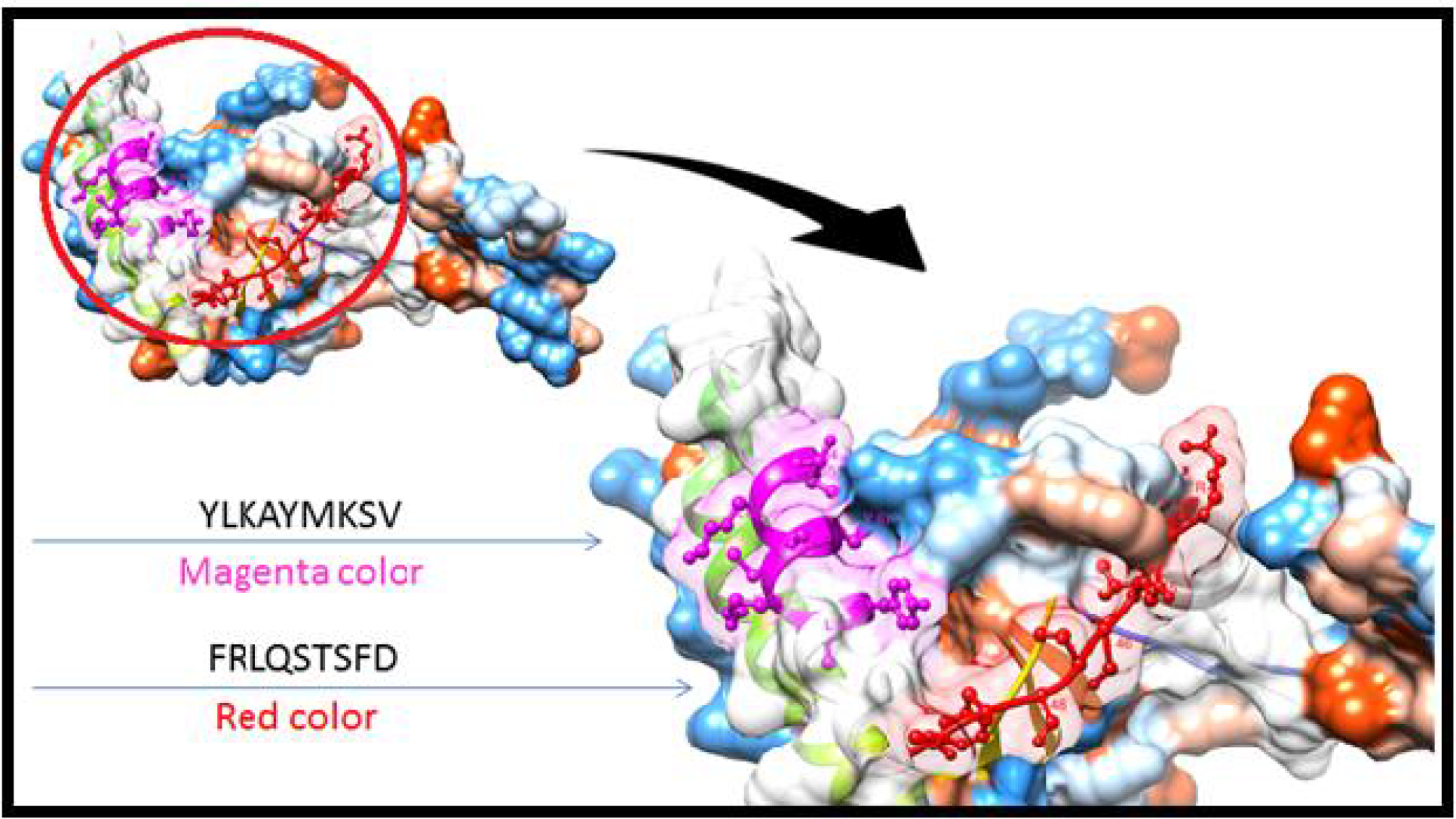
T cell epitopes proposed that interact with MHCII. The arrow show position of(YLKAYMKSV) with magenta color and (FRLQSTSFD) with red in structural level of glycoprotein of *Madurella Mycetomatis TCTP*

**Fig 9.**
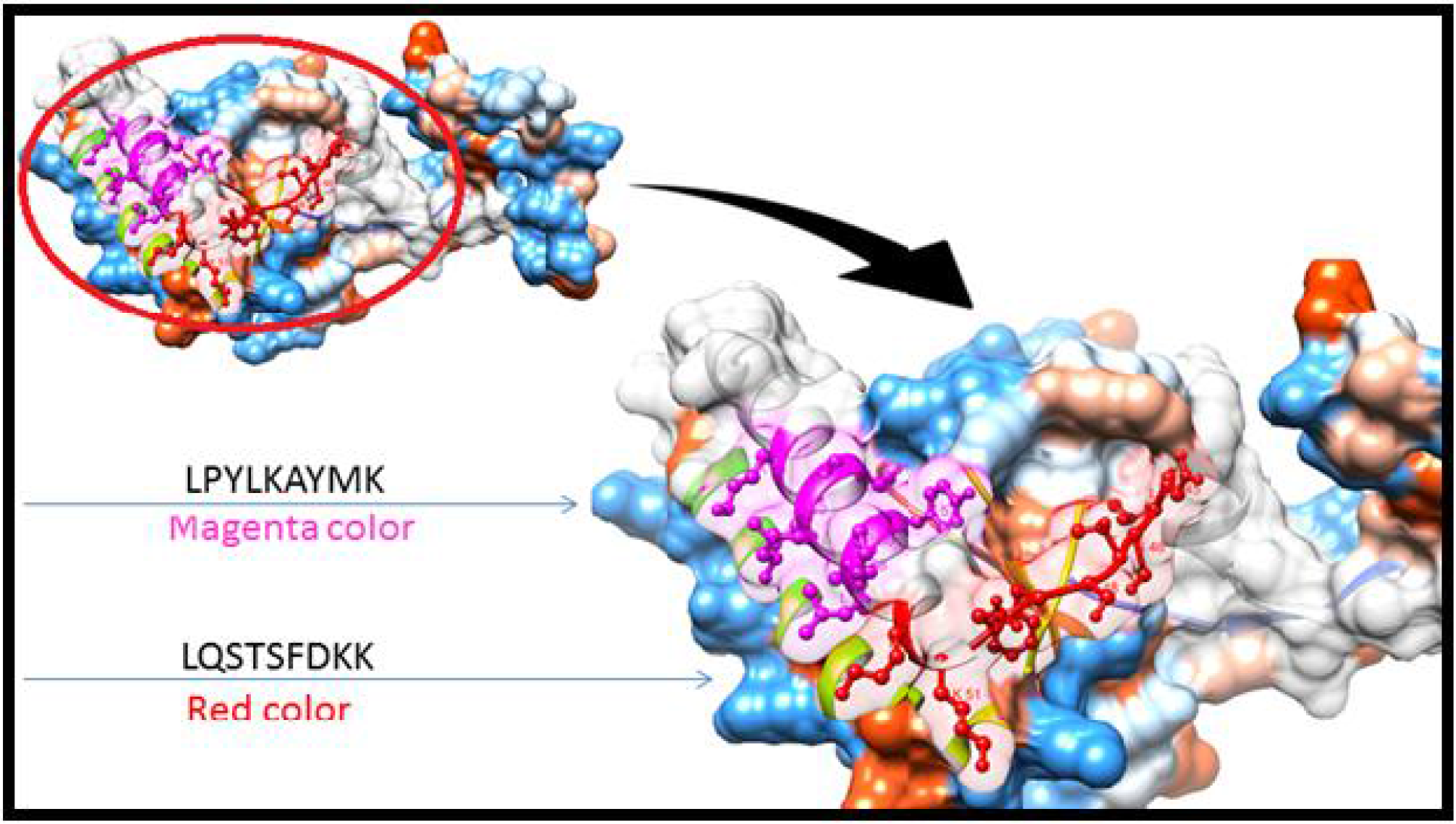
T cell epitopes proposed that interact with MHCII. The arrow show position of (LPYLKAYMK) with magenta and (LQSTSFDKK) with red color in structural level of *Madurella Mycetomatis TCTP*

**Fig 10.**
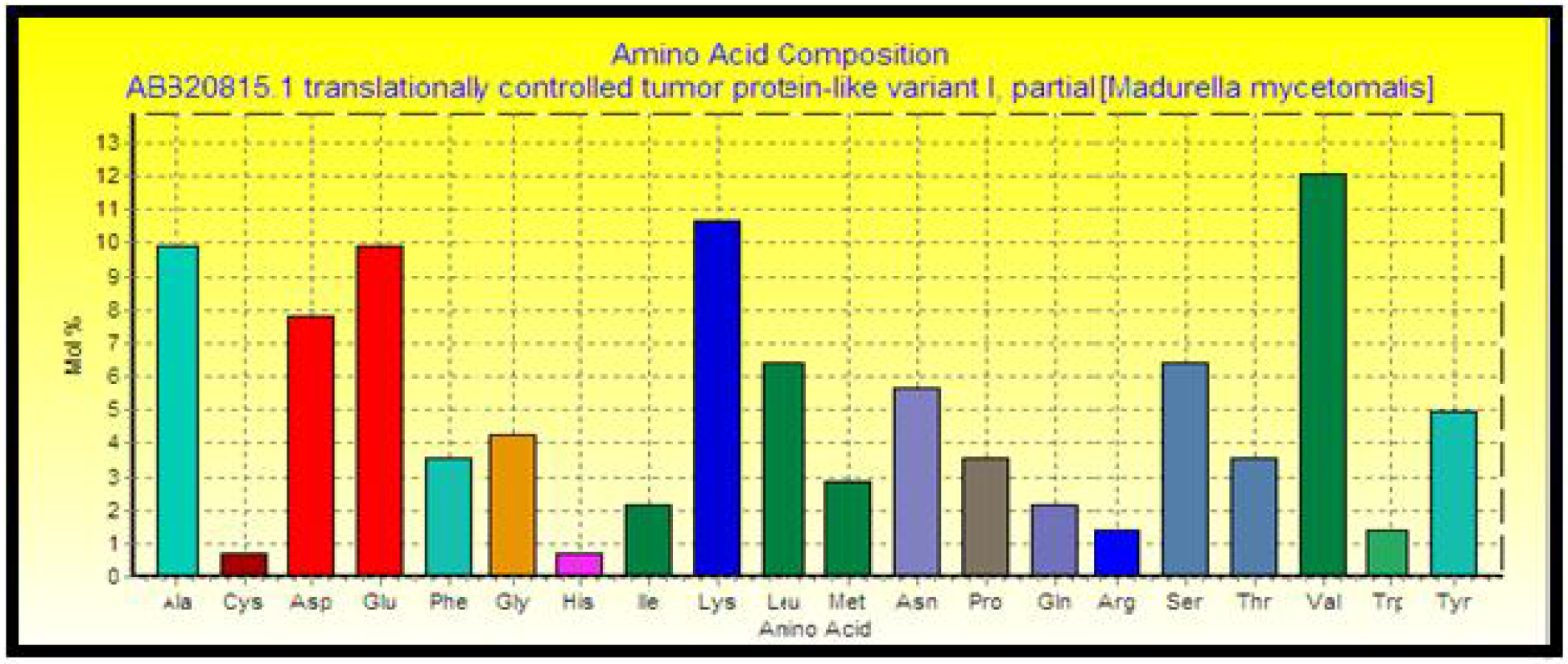
*TCTP* Protein analysis graph (Bioedit program, version 7.0.9.0)

**Table 4.** Atomic composition of *Madurella mycetomatisTCTP*.

| Elements | Symbol | No. of atoms |
| --- | --- | --- |
| Carbon | C | 705 |
| Hydrogen | H | 1095 |
| Nitrogen | N | 177 |
| Oxygen | O | 224 |
| Sulfur | S | 5 |

**Table 5.** The population coverage of Whole world and Sudan for the most promising MHCI, MHCII, MHC I/ II, Shared epitope as MHCI,II epitopes and their alleles.

| Country | MHCI | MHCII | MHCI,II | Shared epitope as MHCI,II |
| --- | --- | --- | --- | --- |
| World | 85.16% | 55.5% | 93.4% | 66.75% |
| Sudan | 90.39% | 39% | 94.14% | 57.15% |

## Discussion

In our study, different peptide vaccines against *TCTP* of *Madurella. mycetomatis* are presented for the first time.

Design of effective vaccine is essential as protection tool against the disease because vaccines and effective treatments of mycetoma are not available till now (12). Peptide vaccines overcome the side effects of conventional vaccines.

*TCTP* is present in form of two alleles. Across the retrieved sequences of *TCTP,* the region from 42-93 is highly conserved. IEDB tools were used. Seven epitopes from the conserved regions were predicted eliciting B lymphocyte using Bepipred test, 37 epitopes were potentially in the surface by passing Emini surface accessability test while 15 epitopes were antigenic using Kolaskar and Tongaonkar test. Only four epitopes passed the three tools (DEVKEFETKAQAYV, EVKEFETKAQAYV, EFETKAQAYV, ETKAQAYV).

111 epitopes were predicted to interact with MHCI alleles with IC50 < 500. Three of them were most promising (YLKAYMKSV, TSFDKKSYL, YMKSVKKAL). 274 predicted epitopes were interacted with MHCII alleles with IC50 < 100. Four of them were most promising (FRLQSTSFD, YLKAYMKSV, YLPYLKAYM, LQSTSFDKK).

Whole world, Sudan and Mexico had the highest population coverages concerning the promising peptides with high affinity to MHC class I alleles alone and the promising peptides with high affinity to MHC class I and class II alleles combined together.

The epitope (YMKSVKKAL) was the most promising one concerning its binding with MHCI alleles, while (FRLQSTSFD) was the most promising for MHC II. The epitope (YLKAYMKSV) is shared between MHC I and II. For the population coverage of *M. mycetomatis TCTP* vaccine, Sudan (90.39%) had the highest percentage for MHC I.

The whole world beside the mycetoma belt will benefit from this vaccine upon its successful development. TCTP is also present in many other species, so predicted peptides can be used as mulifactorial vaccine to protect from many diseases at the same time.

Many studies predicted peptide vaccines for different microorganisms such as, Rubella, Ebola, Dengue, Zika, Lagos rabies virus, HPV, HIV, HCV, malaria, anthrax, influenza, and swine fever using immunoinformatics tools (37–43). A published study found that the peptide FFKEHGVPL had a very strong binding affinity to MHCII alleles which made it a strong candidate as epitope based vaccine against Fructose bisphosphate aldolase of *Madurella mycetomatis* (44). Limitations were seen upon the calcutation of the population coverages concerning the promising peptides with MHC class II for (Venezuela, Senegal, Mexico, India, Colombia and Argentina), beside the combined MHC class I and class II for (Senegal, Mexico, India, Colombia, Argentina and Venezuela)due to missed alleles in the database. The peptide YLKAYMKSV is shared between MHC class I and class II but its percentage in the population coverage was not so high., Further invivo and invitro studies are recommended to prove the effectiveness of these peptides for vaccination.

## Conclusion

We conclude that due to the fact that Peptide vaccines overcome the side effects of conventional vaccines, our study presented for the first time different peptide vaccines against *TCTP* of *Madurella. Mycetomatis.* The epitope (YMKSVKKAL) was the most promising one concerning its binding with MHCI alleles, while (FRLQSTSFD) was the most promising for MHC II. The epitope (YLKAYMKSV) is shared betweenMHC I and II. For the population coverage of *M. Mycetomatis TCTP* vaccine Sudan (90.39%) had the highest percentage for MHC I.

## Acknowledgment

Deep thanks to staff of Africa City of Technology. Pure thanks to everyone who supported us to conduct this paper.

